# An in-silico based clinical insight on the effect of noticeable CD4 conserved residues of SARS-CoV-2 on the CD4-MHC-II interactions

**DOI:** 10.1101/2020.06.19.161802

**Authors:** Selvaa Kumar C, Senthil Arun Kumar, Debjani Dasgupta, Haiyan Wei

**Affiliations:** School of Biotechnology and Bioinformatics, D. Y. Patil Deemed to be University, Sector-15, CBD Belapur. Navi Mumbai-400614, India; Department of Endocrinology and Metabolism, Genetics, Henan children’s hospital (Children’s hospital affiliated to Zhengzhou University), No-33, Longhu Waihuan East road, Zhengzhou-450018, China

**Keywords:** CD4, conserved residues, MHC-II, receptor binding domain, spike S1 protein, SARS-CoV-2

## Abstract

The study is aimed to unveil the conserved residues of CD4 in the context of its purposeful interaction with MHC-II at the receptor-binding domain (RBD) of SARS-CoV-2 compared with the envelope (Env) glycoprotein (gp) 120 of HIV-1. The paired CD4 conserved residues, including the matched CD4 interacting MHC-II epitopes of the structural viral protein domains, were chosen for the protein modelling using the SWISS-MODEL online server. Energy minimization and structural validation of the modelled viral protein domains, including the CD4 and MHC-II protein were achieved by CHIMERA and PROCHECK-Ramachandran Plot respectively. Protein-protein docking was performed by the HADDOCK online tool. The binding affinity score was measured using the PRODIGY online server.

As per our docking report, the Env gp120 of HIV-1 with three identical and three conserved residues of CD4 exhibited the highest binding affinity (−13.9 kcal/mol) with MHC-II than the second-highest RBD-S1-SARS-CoV-2 (−12.5 kcal/mol) with three identical and a single conserved residue of CD4. With a noticeable single salt bridge formation identified at the interacting residues Lys305 (of Env gp120-HIV-1) and Glu139 (of MHC-II); the Env gp120 interaction with MHC-II occupied the crucial His144 and Glu194 (salt-bridge) interacting residues of CD4 with the measured buried surface area 2554.8±40.8 Å^2^. Similarly, the RBD-S1-SARS-CoV-2-MHC-II complex showed two salt bridge formations at the residue sites: 1) Arg567 (of SARS-CoV-2)-Glu194 (of MHC-II) 2) 2) Asp568(of SARS-CoV-2)-Arg165 (of MHC-II) with the increased buried surface area of 1910.9±97.1 Å^2^ over the SARS-CoV score 1708.2±50.8 Å^2^; that camouflaged all crucial CD4 interacting residues of MHC-II. In conclusion, the noticeable conserved residues of CD4 at the RBD-S1 sites of SARS-CoV-2 could interrupt the aspired CD4-MHC-II interactions of adaptive immune activation.

## 1. Introduction

SARS-CoV-2 pandemic has inflicted higher mortality among the global population (https://www.worldometers.info/coronavirus/). Tackling the Severe Respiratory Syndrome Coronaviruses 2 (SARS-CoV-2) human-human transmission chain as well as its diverse virulence characteristics becomes challenging to all the clinicians and researchers worldwide [1]. Its unprecedented spread rate has embraced SARS-CoV-2 to be the dreadful human-threat alike the other deleterious virulent viruses like the Middle East Respiratory Syndrome Coronaviruses (MERS-CoV), Severe Respiratory Syndrome Coronavirus (SARS-CoV) and human immunodeficiency virus type 1 (HIV-1) [2]. Based on the noticeable clinical symptoms of SARS-CoV-2, experts from the global countries have been widely focusing on all potential SARS-CoV-2-based and human-host based therapeutic targets to control its emerging spread rate as well as its virulence among the humans [3–5]. Cytokine storm has been witnessed as one of the major detrimental clinical symptoms of SARS-CoV-2 virulence causing maximum mortality with severe respiratory ailments among the severely ill patients [6–8]. However, with the noticeable decrease in the production of the anti-inflammatory cytokines, especially IL-4, and its concordant CD4+T helper cells counts on the SARS-CoV-2 patients [9–12]. It is more likely for the adaptive immune system to get disturbed mimicking the clinical conditions of the HIV-1 affected patients [12–14].

MHC-II comprised of two domains α1 and β1 derived from the single heavy-α and -β chains that got slightly curved into a β-sheet conformation at the base (for the peptide-antigen binding) with the two α-helices at its top to accommodate the antigenic peptides [15]. The peptide binding unit is further strengthened by the two membrane-proximal immunoglobins (Ig) domains: α2 and β2 adhered to each α1 and β1 heavy chains of MHC-II, respectively [15]. The residues 137-143 of β2-microglobulin domain of MHC-II has been exclusively engaged in interacting with the three exposed loops of the CD4 coreceptor domains D1 and D2 [16,17] (Figure 1). CD4 receptor being the noticeable host-based target of the virulent HIV-1 possessed a conserved binding site: SLWDQ at the L2-loop that exclusively interacts with the antigen-presenting human leukocyte Antigen (HLA)-DP haplotype of the class-II Major histocompatibility complex (MHC-II) expressed by the dendritic cells, macrophages, and B-lymphocytes [18]. Also, the amino acid residues: LNGQEETAGVVSTN (residues length: 170-183) present in the exterior loop of the β1-chain of the MHC-II are engaged in establishing a stable interaction with the CD4 + T-helper cells [18].

**1.**
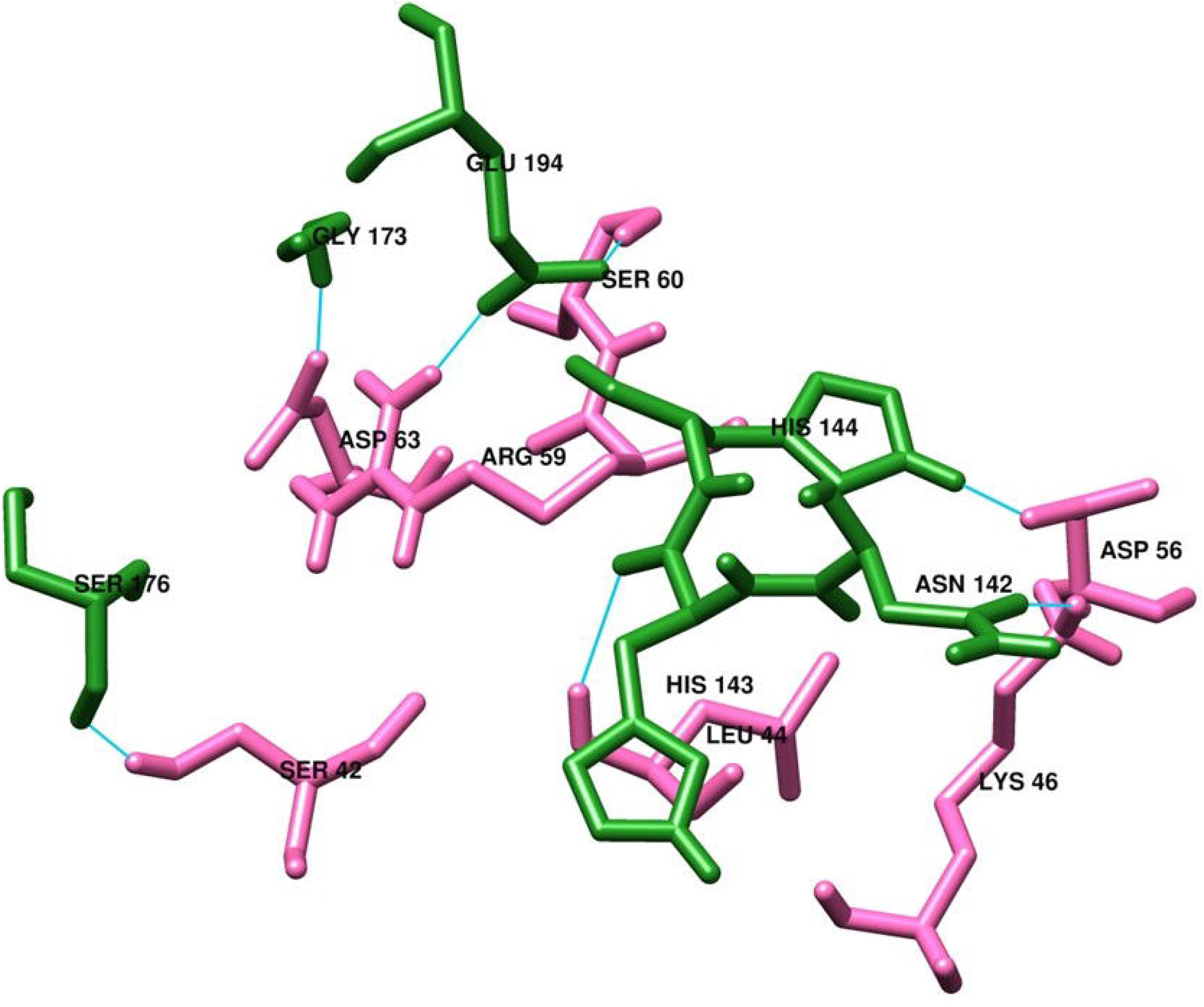
Structural illustration of the CD4-bound MHC-II complex shown in ribbon module. The D1 domain of CD4 comes in proximity with the β domain of MHC-II.

Generally, the HXB2 residues (418-445 residues) spotted at the C-terminal half of the V3 loop of HIV-1 envelope glycoprotein120 (Env gp120) attenuates the adaptive immune response via establishing a stable interaction with the loop 1 (L1) of CD4 [19–21]. Moreover, the HIV-1 Env gp120 by adopting the homology residues of CD4 (SLWDQ) and MHC-II (VVSTQLLLNG), attenuated the CD4-MHC-II interactions and its associated adaptive immune response [18]. Similarly, the homologous residues of IL-2 spotted at the human immune deficiency virus (HIV)-viral receptor proteins were proposed to inhibit the IL-2 mediated immune response [22]. Based on these clinical inferences, we speculate that the receptor-binding domain (RBD) of the spike 1 (S1) protein of SARS-CoV-2 might possess the homologous residues of SLWDQ and/or of any conserved residues of the host-CD4 that seals its interaction with the MHC-II.

To prove this, the prevalence of any conserved CD4 binding residues at the RBD of coronaviruses (SARS-CoV, MERS-CoV, and SARS-CoV-2), and as well as at the Env gp120 (V3 loop) site of HIV-1 were carried out using pairwise sequence alignment. Further, the effect of the existence of any conserved CD4 residues at the RBD sites of coronaviruses in the context of purposeful CD4-MHC-II interactions were studied using the protein-protein docking. The efficacy of the RBD interactions with the MHC-II competing with the CD4 interactions was evaluated based on the Env gp120 (V3 loop) interactions with MHC-II. We believe that SARS-CoV-2 by procuring the crucial conserved binding residues of CD4 with MHC-II could markedly intervene the CD4-MHC-II interactions in the context of adaptive immune activation. Also, all notified CD4 conserved active residues at the RBD sites of SARS-CoV-2 shall be targeted to rejuvenate the disrupted host-adaptive immune response by paving the firm CD4-MHC-II interactions.

## 2. Materials and Methods

### 2.1 Pairwise alignment of the human-CD4 protein sequence with the structural viral protein sequence of coronaviruses and HIV-1

The alignment is intended to examine the prevalence of MHC-II interacting CD4-conserved motif, including the CD4 interacting MHC-II epitopes: SSKRFQPFQQFGRDV and SLWDQ residues in the receptor-binding domain (RBD) of the spike (S1) protein of MERS-CoV, SARS-CoV, and SARS-CoV-2 [18,23]. Concomitantly, the study has employed the CD4-SLWDQ comprised envelope glycoprotein 120 of HIV-1 (Env gp120-HIV-1) as the guided protein study model to fulfil the objective of the study. Protein sequences of CD4 (Accession number:P01730), MHC-II (Accession number:U3PYM0), and Env gp120-HIV-1 (Accession number:P04582; residues 33-506) were downloaded from the Uniprot database [24]. Alternatively, the protein sequences of SARS-CoV-1 surface glycoprotein (Accession number:NP_828851.1), MERS-CoV (Accession number:AYN72346.1), and SARS-CoV-2 (Accession number:QHD43416.1) were downloaded from the NCBI database (https://www.ncbi.nlm.nih.gov/) To identify the SLWDQ conserved motif and its flanking residues, including the active conserved residues of the CD4 co-receptor in the sequence of the above-mentioned virulent viruses; CD4 of the *host-Homo sapiens* was aligned with each of these protein sequences: 1) Env gp120 of HIV-1 2) SARS-CoV-1 3) SARS-CoV-2 and 4)MERS-CoV using Clustal Omega online server [25]. Amino acids in concordant with the better alignment report were chosen for generating the protein 3D structures.

### 2.2 Homology modelling of the human-CD4 and MHC-II protein templates, and the structural viral protein domains of SARS-CoV family and HIV-1

The entire downloaded protein sequences of RBD of the S1 protein of coronaviruses (SARS-CoV, MERS-CoV, and SARS-CoV-2) and Env gp120 of HIV-1, including the host-human based MHC-II and CD4 protein sequences were chosen for the homology modelling using the SWISS-MODEL online server [26]. By referring to the enlisted templates, the overall resolution, query coverage and the residue identity were chosen for the homology modelling. All structurally modelled protein structures were subjected to energy minimization followed by its structural validation using CHIMERA [27] and ProSa-Web online server [28], respectively. Parallelly, all generated protein models were structurally examined by validating its phi-psi plot using the PROCHECK-Ramachandran plot [29].

### 2.3 Protein-protein docking of the structural viral protein domains with MHC-II in compliance to the CD4-MHC-II interactions

Protein-protein docking was performed using the modelled protein structures in the sequential order by the HADDOCK (High Ambiguity Driven protein-protein DOCKing) online server [30]. HADDOCK has been classified as the information-oriented flexible docking module. In the context of the evaluation of MHC-II-SARS-CoV-2 (antigen) presentation, initially, the protein-protein docking was performed between the human-MHC-II protein chain B (H-2 CLASS II Histocompatibility Antigen) and human-CD4 coreceptor. Upon this MHC-II-CD4 docking, the active residues engaged in establishing the stable interactions were chosen in each of the MHC-II and CD4 binding domains. The active residues spotted at the CD4 interacting loci includes Arg35, Tyr40, Phe43, Leu44, Trp45, Lys46, Arg59, Ser60 and Asp63. While, the active residues like Leu114, Val116, Val142, Val143, Ser144, Thr145, Ile148, Leu158, Met160 and Glu162 were spotted at the MHC-II interacting loci. Moreover, these active site residues of each MHC-II and CD4 interacting loci, respectively, were procured by the literature review [31]. By default, all passive residues that are generally classified as the surrounding surface residue would be defined and marked by the software on its own. Following the protein-protein docking of each of the RBD of the S1 protein of coronaviruses (SARS-CoV, MERS-CoV, and SARS-CoV-2) and the Env gp120 of HIV-1 with MHC-II was performed sequentially. The HADDOCK score and the buried interface regions were calculated during docking. Since the calculated HADDOCK scores don’t have any unit values as such, we have chosen the PRODIGY online server [32] to measure the binding affinity in *kcal/mol.*

## 3.0 Results

### 3.1 Pairwise alignment of the human-CD4 with the V3 loop of Env gp120-HIV-1 and the RBD of the S1 protein of coronaviruses

As evidenced, the crucial CD4-SLWDQ residue has been spotted as the partial segment of the major interface region with the residues Arg35, Tyr40, Phe43, Leu44, Trp45, Lys46, Arg59, Ser60 and Asp63 that profusely strengthens the CD4 interactions with MHC-II [33]. Thus, the entire MHC-II interacting residues from Arg35-Asp63 (residues count 29) of CD4, including the crucial short-SLWDQ residue were chosen for the pairwise alignment. Firstly, to locate the short CD4-SLWDQ residue accompanied with its other flanking (Arg35-Asp63) residues in the V3 loop of Env gp120-HIV-1; the pairwise alignment was performed between the host-human CD4 and the V3 loop of Env gp120-HIV-1. Noticeably, three identical residues with four conservative substitutions with three marked gaps were spotted with the CD4-Env gp120-HIV-1 alignment. Secondly, upon the pairwise alignment performed between the human-CD4 and the RBD of the S1 protein of SARS-CoV, its neither the identical residues nor the conservative substitutions were spotted with 14 gaps in total. The overall gaps have extended the length of the alignment between SARS-CoV and CD4. Thus, in comparison with SARS-CoV-2 wherein the alignment ends with ‘VRDP’ there is an additional thirteen amino acids observed between them. As a result, we were compelled to trim the size of the alignment matching with that of SARS-CoV2 (termed as *trimmed)* while keeping the complete stretch of alignment as such (termed as *complete)* In case of the CD4-MERS-CoV pairwise alignment, it’s only a single identical residue was spotted with the noticeable 24 gaps. Thirdly, the pairwise alignment of the CD4-SARS-CoV-2 showed three identical residues with a single conservative substitution and a noticeable gap (Figure 2a-d).

**2.**
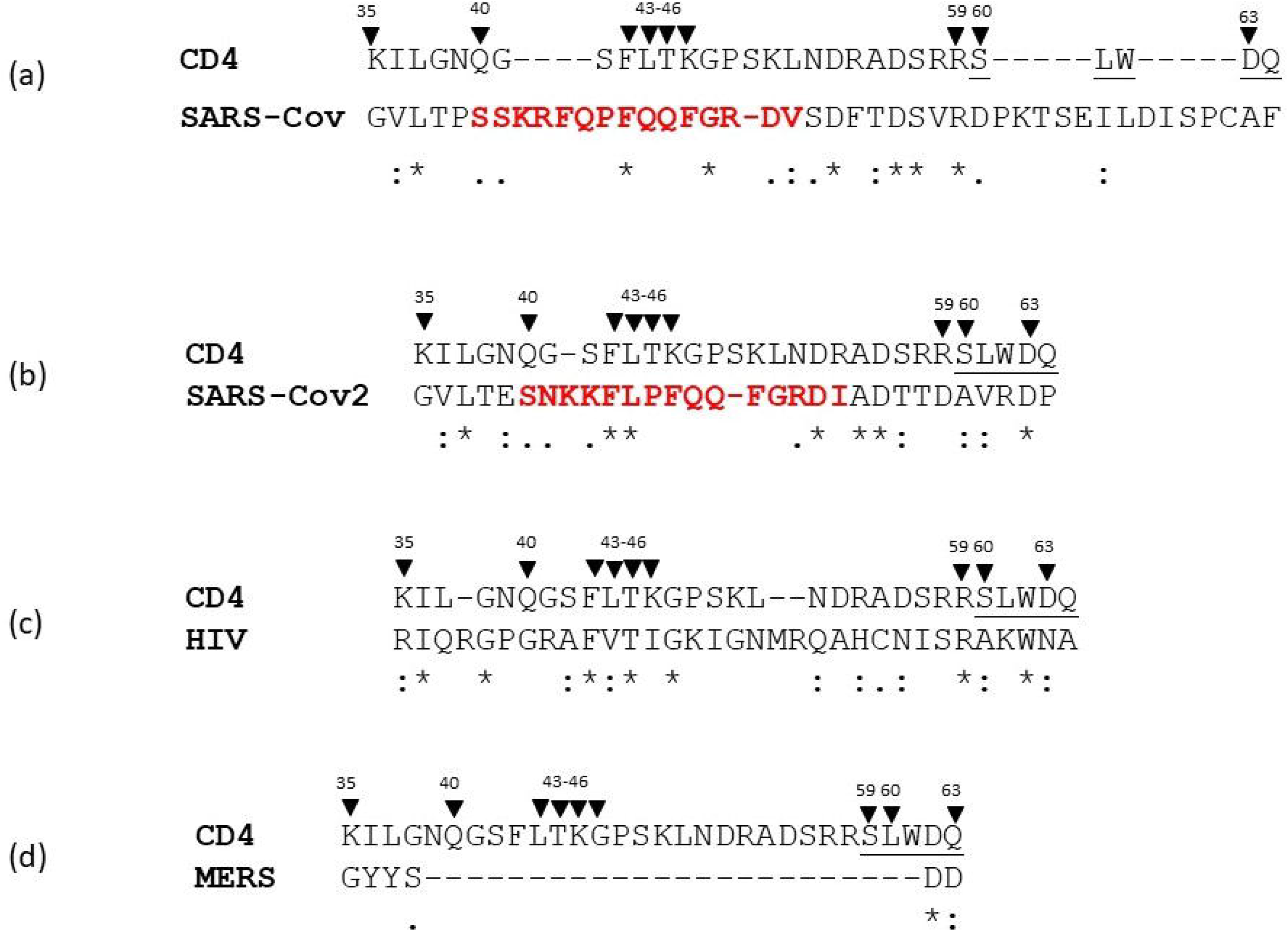
Pairwise sequence alignment of the MHC-II interacting CD4 protein sequence with the structural receptor-binding domain (RBD) of the spike 1 (S1) protein of coronaviruses (SARS-CoV, SARS-CoV-2, and MERS-CoV) and the envelope (Env) glycoprotein (gp) 120 of HIV-1. The underlined short nucleotide motif SLWDQ is linked with the immunosuppression. Exome highlighted in red font implies the MHC-II conserved epitopes. All noticeable amino acid residues of the above-mentioned structural viral proteins, including the human-CD4 domain residues that are indulged in stable interactions with MHC-II, have been highlighted using downward black arrows (). Homology residues spotted with the paired nucleotides have been marked using an asterisk (*) symbol. Protein sequences with conservative substitutions have been highlighted using a semicolon (:). Amino acids with weak substitutions have been spotted by a single dot (.)

### 3.2 Homology modelling of the structural viral membrane proteins and the host human-CD4 and -MHC-II proteins

The protein sequences of CD4, MHC II, Env gp120-Hiv, SARS-CoV and SARS-CoV-2 were chosen for model building respectively. The most preferred template for 3D modelling of Env gp120-HIV-1 was PDB ID: 3J70 [34]) with 97.01% sequence identity and 97% query coverage; PDB ID: 6acd [35]) with 99.925 % sequence identity and 95% query coverage for SARS-CoV (RBD of S1); PDB ID: 6m18 chain B [36]) with 100 % sequence identity and 99% query coverage for SARS-CoV-2 (RBD of S1); PDB ID: 1JL4 [37]) with missing residues for human-CD4; PDB ID: 4Z7U [38] with 100% sequence identity and 74% query coverage for human-MHC-II. All chosen crystal structures with their respective PDB ID’s were considered for model building using SWISS-MODEL online server to generate their respective 3D model structures. The generated 3D model structure of each of the study proteins was employed for its energy minimization by CHIMERA followed by its structural validation by ProSA-Web and Ramachandran Plot analysis. The ProSA-web analysis for all the 3D-modelled study proteins showed a better-localized model quality to perform the protein-protein docking (Supplementary material 1a). While with the Ramachandran plot analysis, the following structural conformations has been witnessed with each of the 3D-modelled study proteins such as the human-CD4 protein model showed 74% residues in the highly favoured regions, 24% in the additional allowed regions, 1.3% in the generously allowed regions, and 1.3% in the disallowed regions; the Env gp120-HIV-1 protein model showed 89.2% residues in the highly favoured regions, 7.4% in the additional allowed regions, 1.4% in the generously allowed regions, and 1.9% in the disallowed regions; the MHC-II protein model showed 93.6 % residues in the highly favoured regions, 5.3% in the additional allowed regions, 0.6% in the generously allowed regions, and 0.6% in the disallowed regions; the SARS-CoV-2 protein model showed 86.1% residues in the highly favoured regions, 11.9% in the additional allowed regions, 1.7% in the generously permissible regions, and 0.3% in the disallowed regions; the SARS-CoV protein model showed 78.1 % residues in highly favoured regions, 19.9% in the additional allowed regions, 1.6% in generously allowed regions, and 0.3% in the disallowed regions (Supplementary material 1b).

### 3.3 Protein-protein docking of structural viral membrane proteins and human-CD4 with the MHC-II

The HADDOCK score generated upon MHC-II-CD4 docking was lower (−100.7±-2.9) than the HADDOCK score value of MHC-II-Env gp120-HIV-1 complex (−111.1±-3.8) and the RBD-SARS-CoV-2 complex (−122.1±-5.0). The HADDOCK score value of the MHC-II-RBD-SARS-CoV complex was the least amongst all the structural viral study proteins with the score value of −55.7±-5.5 (in the *complete* state) and −84.0±-2.6 (in the *trimmed* state). Nevertheless, of all the studied proteins, the highest HADDOCK score value was recorded with the MHC-II-RBD-S1-SARS-CoV-2 complex. Commonly, the HADDOCK score value is the culmination of the energies measured by the OPLS force field-based electrostatics and the *Van der* walls accompanied by the evaluation of the buried surface area and desolvation.

It is quite obvious that a firm conclusion on the docking results could have arrived only after the comparative analysis of all diverse data generated for each study protein complexes. For the same reason, we have adopted the PRODIGY online server tool to estimate the binding affinity of all the docked protein complexes. The PRODIGY analysis showed the least binding affinity score value (−9.4 kcal/mol) of the MHC-II-CD4 protein complex. While the MHC-II-Env gp120-HIV-1 complex showed the highest binding affinity (−13.9 kcal/mol) followed by the MHC-II-RBD-S1-SARS-CoV-2 complex (−12.5 kcal/mol) and the MHC-II-RBD-S1-SARS-CoV ((−11.9 kcal/mol (in the *complete* state) and −11.8 kcal/mol (in the *trimmed* state)).

In terms of the buried surface area, the MHC-II-CD4 complex has recorded the least score (1183.6 ±72.7 Å^2^), while, the MHC-II-SARS-CoV (in the *complete* state) showed the highest score (2710.3 ±104.8 Å^2^) of all the experimental study proteins. The MHC-II-HIV-1 complex recorded the second-highest buried surface area score value (2554.8 ±40.8 Å^2^) followed by the MHC-II-SARS-CoV-2 complex (1910.9 ±97.1 Å^2^) and the MHC-II-SARS-CoV complex (1708.2 ±50.8 Å^2^ in the *trimmed* state). Among the coronaviruses, we still consider the MHC-II-SARS-CoV-2 complex to possess the highest buried surface area score value over the MHC-II-SARS-CoV complex score (in the *complete* state) as the score value gets influenced by the supplementary residues as witnessed in the pairwise alignment (Figure 3a-d) contributing a larger buried surface area to RBD-S1-SARS-CoV than its counterpart (Table 1).

**3.**
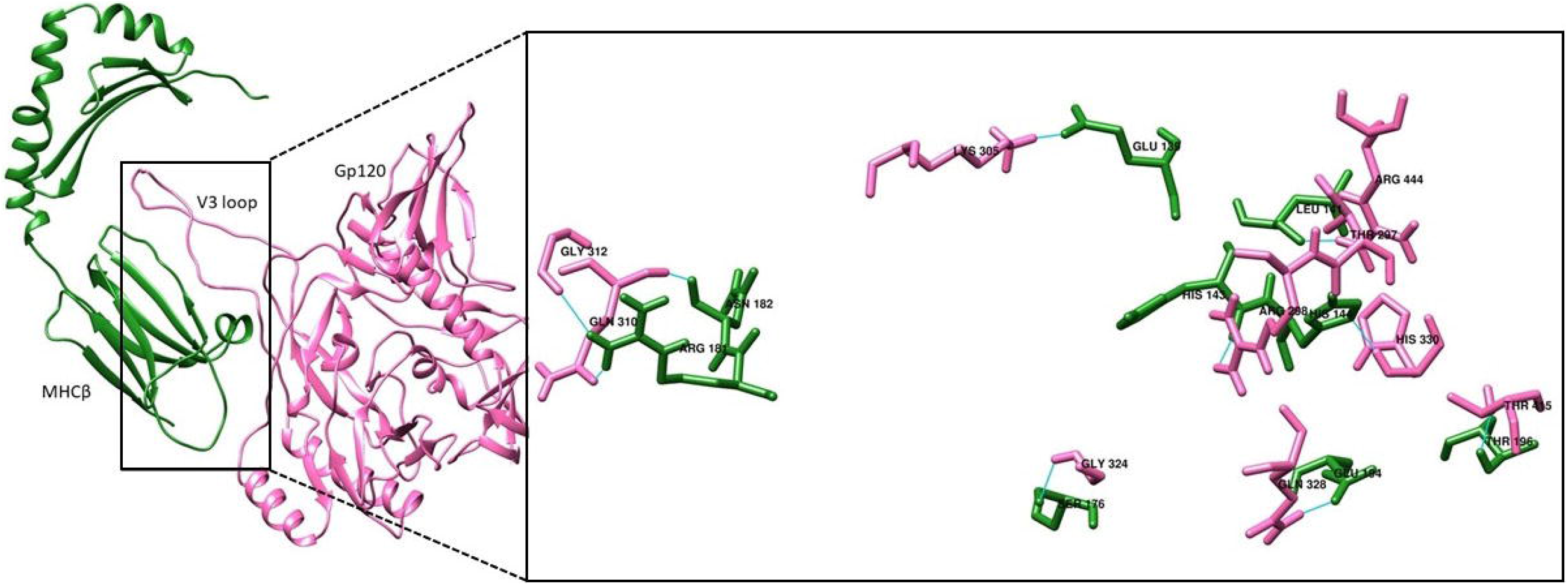
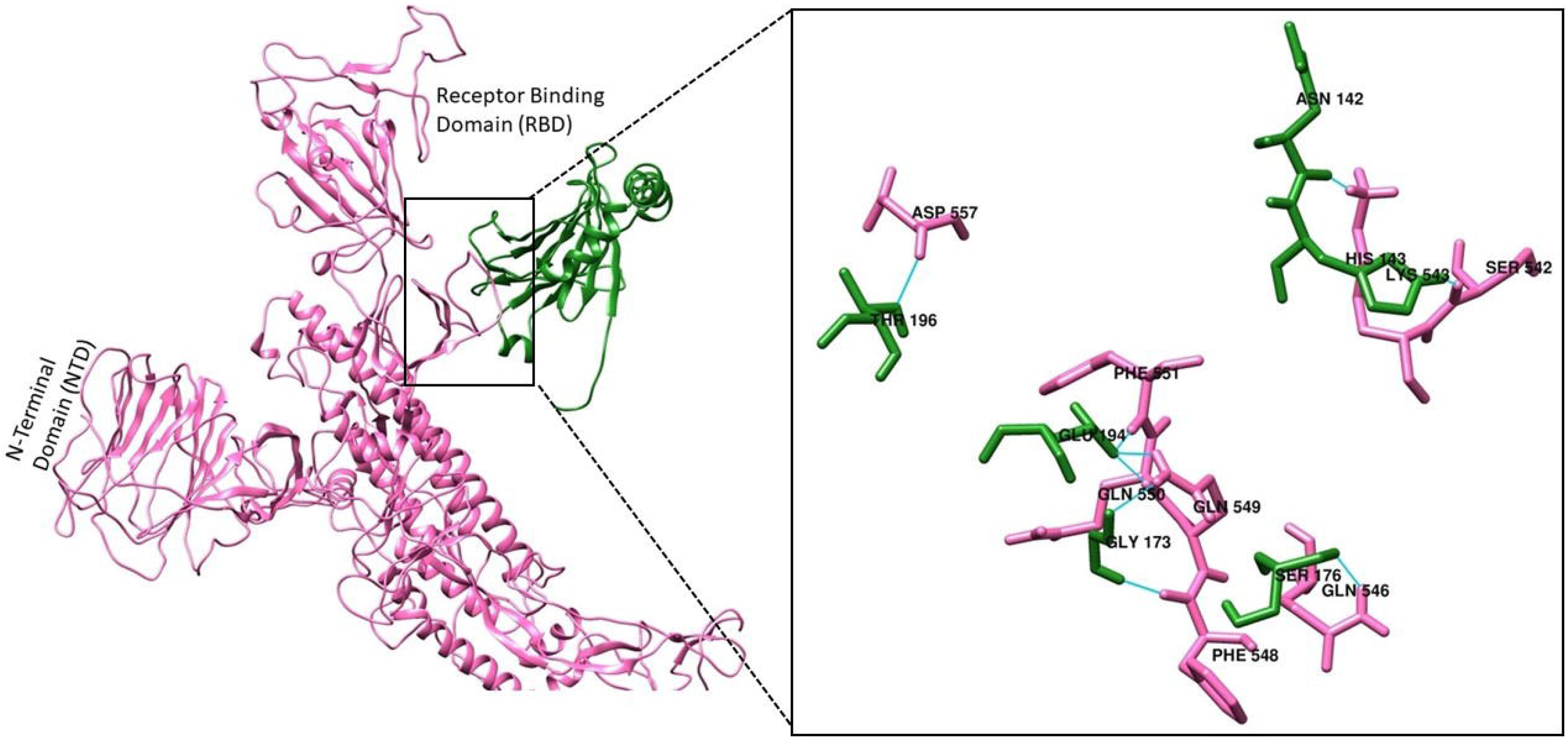
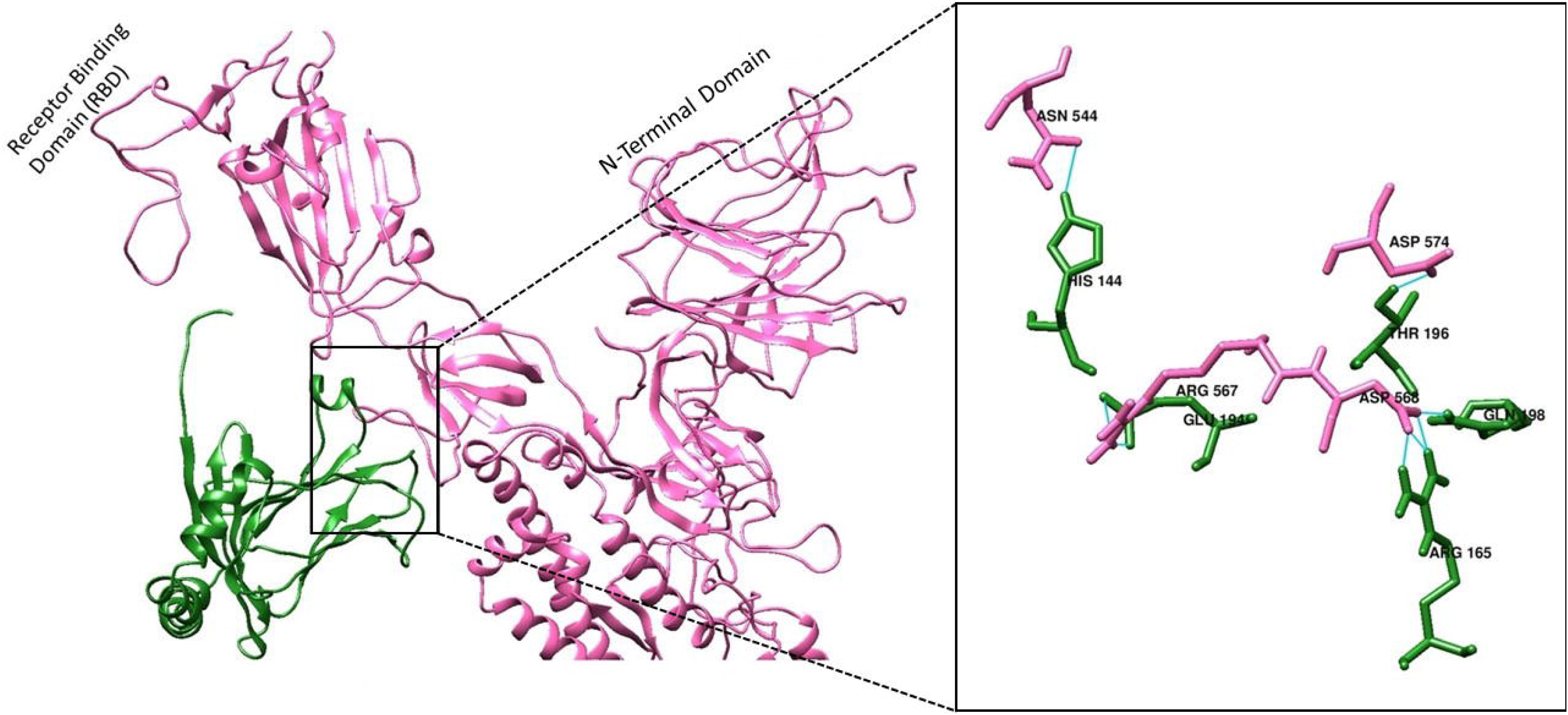
Protein-protein docking of the structural viral membrane proteins with the CD4. (a) Structural representation of the envelope (Env) glycoprotein (gp) 120 (V3 loop of HIV-1) interactions with the CD4; showing the V3 loop with the CD4 homology residues established direct contact with the MHC-II-β domain. (b) Structural protein docking of the receptor-binding domain (RBD) of SARS-CoV with MHC-II showed the RBD tail end (the MHC-II epitope) established direct contact with the MHC-II-β domain. (c) Structural protein docking of the RBD of SARS-CoV-2 with MHC-II showed the RBD tail end (the MHC-II epitope) established direct contact with the MHC-II-β domain. (Note: All noticeable amino acid interactions have been shown in the stick file format).

**Table 1.** Protein-Protein docking using HADDOCK software and the binding affinity calculation using PRODIGY software

| Protein-complex | HADDOCK values |  | PRODIGY values |
| --- | --- | --- | --- |
| | HADDOCK score | Buried surface Area ($\text{\AA}^2$) | |
| MHC II -CD4 | -100.7 +/- 2.9 | 1183.6 +/- 72.7 | -9.4 kcal/mol |
| MHC II-HIV-1 | -111.1 +/-3.8 | 2554.8 +/- 40.8 | -13.9 kcal/mol |
| MHC II-SARS-cov(Complete) | -55.7 +/-5.5 | 2710.3 +/- 104.8 | -11.9 kcal/mol |
| MHC II-SARS-cov(trimmed) | -84.0 +/- 2.6 | 1708.2 +/- 50.8 | -11.8 kcal/mol |
| MHC II-SARS-Cov-2 | -122.1 +/-5.0 | 1910.9 +/- 97.1 | -12.5 kcal/mol |

In the context of interaction module, the MHC-II-CD4 complex exhibited two salt bridges: one with His144 (of MHC-II) and Asp56 (of CD4), and another with Glu194 (of MHC-II) and Arg59 (of CD4) supported with five hydrogen bonds (Table 2) (Figure 2a). The Env gp120 of HIV-1 developed a single salt bridge with Glu139 residue (of MHC-II) interacting with the Lys305 residue (of Env gp120-HIV-1), which indeed has been supported with ten hydrogen bonding residues, including the CD4 interacting His144 and Glu194 (salt-bridge forming) residues of MHC-II (Figure 2b). The MHC-II-RBD-S1-SARS-CoV complex didn’t show any salt bridge conformations between the residues, nonetheless, about seven hydrogen bonding residues were witnessed, including the CD4-interacting Glu194 (saltbridge forming) residue of MHC-II. Of the seven major CD4 interacting MHC-II residues (marked with an asterisk (*)), five mapped residues (marked with *), including the His144 and Glu194 (of MHC-II) were indulged in hydrogen bonding with Env gp120-HIV-1(Figure 2c). Concordantly, the MHC-II-RBD-S1-SARS-CoV-2 complex developed one salt bridge between the CD4 interacting Glu194 residue site (of MHC-II) and the Arg567 residue (of SARS-CoV-2), but the second salt bridge was spotted at the different residue site of Arg165 (of MHC-II) interacting with Asp 568 (of SARS-CoV-2). These two charged amino acids viz. Arg567 and Asp568 from RBD domain of SARS-CoV2 is a segment of MHC-II epitope binding site. Concertedly, these two salt bridges of MHC-II-RBD-SARS-CoV-2 complex has been supported by three hydrogen bonding residues, including, the CD4 interacting His144 (salt bridge forming) residues of MHC-II. While, with the RBD-SARS-CoV-2-MHC-II complex, its only one CD4 interacting mapped residue Glu194 (marked in *) were engaged in the salt bridge formation, however, the other mapped His144 residue (marked in *) engaged in hydrogen bonding (Figure 2d). In case of the MHC-II-RBD-SARS-CoV complex, all three CD4 interacting mapped residues of MHC-II such as Ser176, Glu194 and His143 (marked in *) were engaged in hydrogen bonding without any salt-bridge formation (Table 2).

**Table 2.** The interacting residues between MHC II and the viral proteins reported along with the type of interactions.

| MHC II | CD4 | Type of interaction |
| --- | --- | --- |
| **His144\*** | **Asp56** | Salt-bridge |
| Gly173 | Asp63 | Hydrogen bond |
| Ser176* | Ser42 | Hydrogen bond |
| His143* | Leu44 | Hydrogen bond |
| Asn142 | Lys46 | Hydrogen bond |
| **Glu194\*** | **Arg59** | Salt-bridge |
| Glu194* | Ser60 | Hydrogen bond |
| **MHC II** | **HIV** |  |
| His144* | Thr297 | Hydrogen bond |
| Arg181 | Gln310 | Hydrogen bond |
| Arg181 | Gly312 | Hydrogen bond |
| Thr196 | Thr415 | Hydrogen bond |
| His143* | Arg298 | Hydrogen bond |
| **Glu139** | **Lys305** | Salt-bridge |
| Asn182 | Gln310 | Hydrogen bond |
| Ser176* | Gly324 | Hydrogen bond |
| Glu194* | Gln328 | Hydrogen bond |
| His144* | His330 | Hydrogen bond |
| Leu141 | Arg444 | Hydrogen bond |
| **MHC II** | **SARS-Cov** |  |
| Asn142 | Lys543 | Hydrogen bond |
| Ser176* | Gln546 | Hydrogen bond |
| Gly173 | Gln549 | Hydrogen bond |
| Glu194* | Gln549 | Hydrogen bond |
| Glu194* | Gln550 | Hydrogen bond |
| Glu194* | Phe551 | Hydrogen bond |
| Thr196 | Asp557 | Hydrogen bond |
| His143* | Ser542 | Hydrogen bond |
| Gly173 | Phe548 | Hydrogen bond |
| **MHC II** | **SARS-Cov2** |  |
| **Glu194\*** | **Arg567** | Salt-bridge |
| His144* | Asn544 | Hydrogen bond |
| **Arg165** | **Asp568** | Salt-bridge |
| Thr196 | Asp574 | Hydrogen bond |
| Gln198 | Asp568 | Hydrogen bond |

## 4.0 Discussion

SARS-CoV-2 pandemic has emerged as a dreadful human threat with the increasing number of casualties of the death toll of 430, 133 (as of 14^th^ June 2020) across the global countries (https://www.worldometers.info/coronavirus/). SARS-CoV-2 being remarkably contagious than the highly virulent MERS-CoV and SARS-CoV strains, clinicians have been struggling to either control its spread rate or its virulence among the global population [39,40]. Clinical therapeutic targets of either host-based or the direct SARS-CoV-2-based, especially its spike1 glycoprotein (S1)-receptor binding domain (RBD) have been widely pronounced for the development of vaccines and/or other natural therapeutic bio-actives against the SARS-CoV-2 virulence [41,42]. However, clinicians are yet struggling to find any promising biological targets because of the diverse clinical characteristics of SARS-CoV-2 [43,44]. Of all noticeable clinical symptoms, the cytokine storm and the diminished CD4+T helper cells production have been remarkably noticed among the infected patients [12,45]. Moreover, therapeutic targets in the context of the adaptive immune response regulated by the MHC-II antigen presentation of SARS-CoV-2 remains unexplored demanding more research studies in this clinical perspective [45,46]. We believe that any modules of clinical targets linked with the adaptive immunity to be crucial to tackle the emerging spread rate as well as the pathogenesis of SARS-CoV-2 [46].

Having enlightened on a diverse range of epitopes at the S1-RBD site of SARS-CoV-2 that could indulge in stable interactions with the immune cells such as T-cell, B-cell and MHC-II [47–49]; a noticeable adaptive immune response must have been witnessed among the SARS-CoV-2 infected groups. Contradictorily, a reduced level of CD4+T-helper cell fractions was noticed among the larger group of SARS-CoV-2 patients [9,12,45]. However, the cytotoxic CD8+T-cells levels remain fluctuated among the patient groups within which its hyperactive state might contribute to the severe respiratory ailments associated with the hyper-immune activation and cytokine storm [12,13,50]. This clinical observation has preceded the hypothesis that SARS-CoV-2 might adopt the virulence characteristics of HIV-1 in disrupting the adaptive immune responses initiated by the MHC-II antigen presentation to CD4+T-helper cells. Also, we firmly believe that the prevalence of the conserved active site (surface interface residues) residues of CD4 co-receptor at the S1-RBD site of SARS-CoV-2 could disturb the adaptive immune system by intervening the CD4-MHC-II interactions.

While seeking the prevalence of any CD4 conserved residues (Arg35, Tyr40, Phe43, Leu44, Trp45, Lys46, Arg59, Ser60 and Asp63), including the CD4 homology residues of SLWDQ, and its MHC-II interacting epitopes SSKRFQPFQQFGRDV at the RBD of SARS-CoV viruses(Figure 2a-d) [18,23]; We found the procurement of CD4 conserved residues such as Arg567 and Asp568 accompanied with the CD4 interacting MHC-II epitopes (SNKKFLPFQQ-FGRDI) by the RBD-SARS-CoV-2 has facilitated the stable salt bridge formations with the Glu194 and Arg165 residues of MHC-II, respectively (Table 2). Following, the Env gp120-HIV-1 has recorded the highest binding affinity score for MHC-II amongst all viral study proteins. However, the human-CD4 showed the least binding affinity score for MHC-II with the two noticeable salt bridge formations. The Env gp120-HIV-1 has established a single salt bridge formation with the MHC-II. With this interaction, the entire CD4 interacting residues (marked using *) of MHC-II has been occupied with the Env gp120-HIV-1 binding with the highest buried surface area. Similarly, the RBD-SARS-CoV-2 binding with MHC-II has enclaved its crucial CD4 interacting residues such as His144 and Glu194 with the second-highest buried surface area. Moreover, the RBD-SARS-CoV binding has accommodated all crucial CD4 interacting residues with the MHC-II (marked in *), showed a higher binding affinity and the buried surface area than the CD4-MHC-II complex.

Combinedly, the RBD-SARS-CoV-2 that possessed three identical residues with one conserved residue of CD4 has established a stable interaction with the MHC-II akin to the Env gp120-HIV-1 with three identical residues and three conserved residues. However, the RBD-SARS-CoV by procuring the CD4-interacting MHC-II epitopes: SSKRFQPFQQFGR-DV established a strong interaction with MHC-II compared with human CD4-MHC-II interactions, but showed a weak interaction compared with the HIV-1 and SARS-CoV-2 (Figure 2a, Table 1). This may be due to the absence of any noticeable CD4 conserved residues at the RBD-SARS-CoV. In conclusion, SARS-CoV-2 showing an increased competence in binding with MHC-II, particularly at the crucial CD4 interacting residues could interrupt the purposeful CD4-MHC-II interactions, markedly affecting the MHC-II antigen presentation to CD4+T-helper cells. Indeed, this interruption could efface the development of adaptive immune response with the diminished secretions of CD4+T-helper cells that have been witnessed widely among the severe clinical cases of SARS-CoV-2. Vaccines or monoclonal antibodies targeting these proposed conserved and/or the identical CD4 epitopes of the RBD-SARS-CoV-2 would help to defend the SARS-CoV-2 pathogenesis by restoring the crucial CD4-MHC-II interactions of antigen presentation and its associated adaptive immune response.

## Acknowledgement

The authors would like to acknowledge technical support from the School of Biotechnology and Bioinformatics, D.Y. Patil Deemed to be University, CBD Belapur for providing access to the softwares for biological data analysis. This work was not funded by any external funding agencies.

