## Supplementary material for "An in-silico based clinical insight on the effect of noticeable CD4 conserved residues of SARS-CoV-2 on the CD4-MHC-II interactions": supplemetary material 1(a)Ramachandran plot sars-cov2.pdf

PROCHECK

### Ramachandran Plot

## 5478370

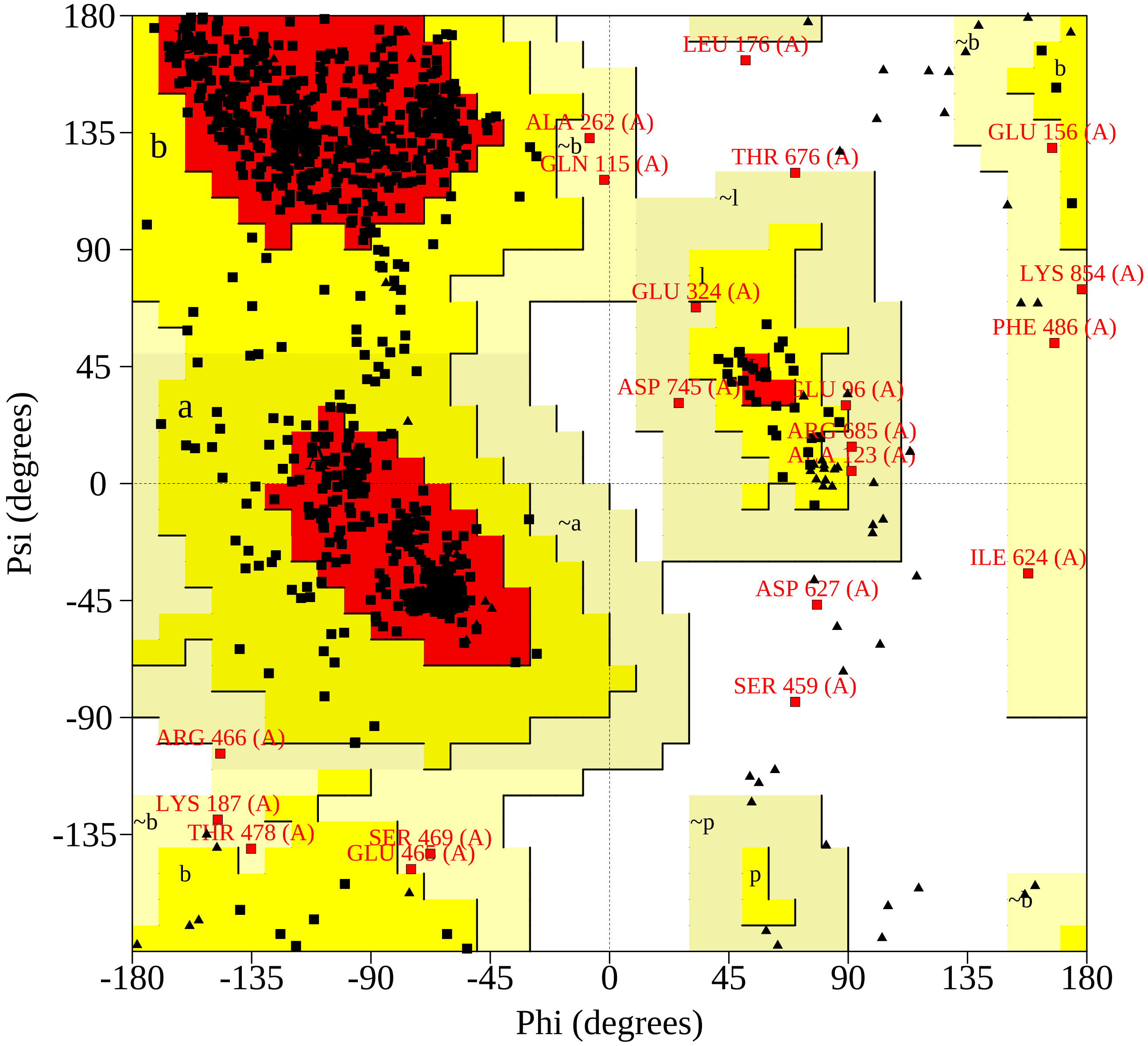


Plot statistics

| | | |
|---|---|---|
| Residues in most favoured regions [A,B,L] | 854 | 86.1% |
| Residues in additional allowed regions [a,b,l,p] | 118 | 11.9% |
| Residues in generously allowed regions [~a,~b,~l,~p] | 17 | 1.7% |
| Residues in disallowed regions | 3 | 0.3% |
|  | ---- | ------ |
| Number of non-glycine and non-proline residues | 992 | 100.0% |
| Number of end-residues (excl. Gly and Pro) | 2 | |
| Number of glycine residues (shown as triangles) | 74 | |
| Number of proline residues | 52 | |
|  | ---- |  |
| Total number of residues | 1120 | |

Based on an analysis of 118 structures of resolution of at least 2.0 Angstroms and R-factor no greater than 20%, a good quality model would be expected to have over 90% in the most favoured regions.

5478370_01.ps
