## Supplementary figures and images for "An in-silico based clinical insight on the effect of noticeable CD4 conserved residues of SARS-CoV-2 on the CD4-MHC-II interactions"

### supplementary material 1b.jpg

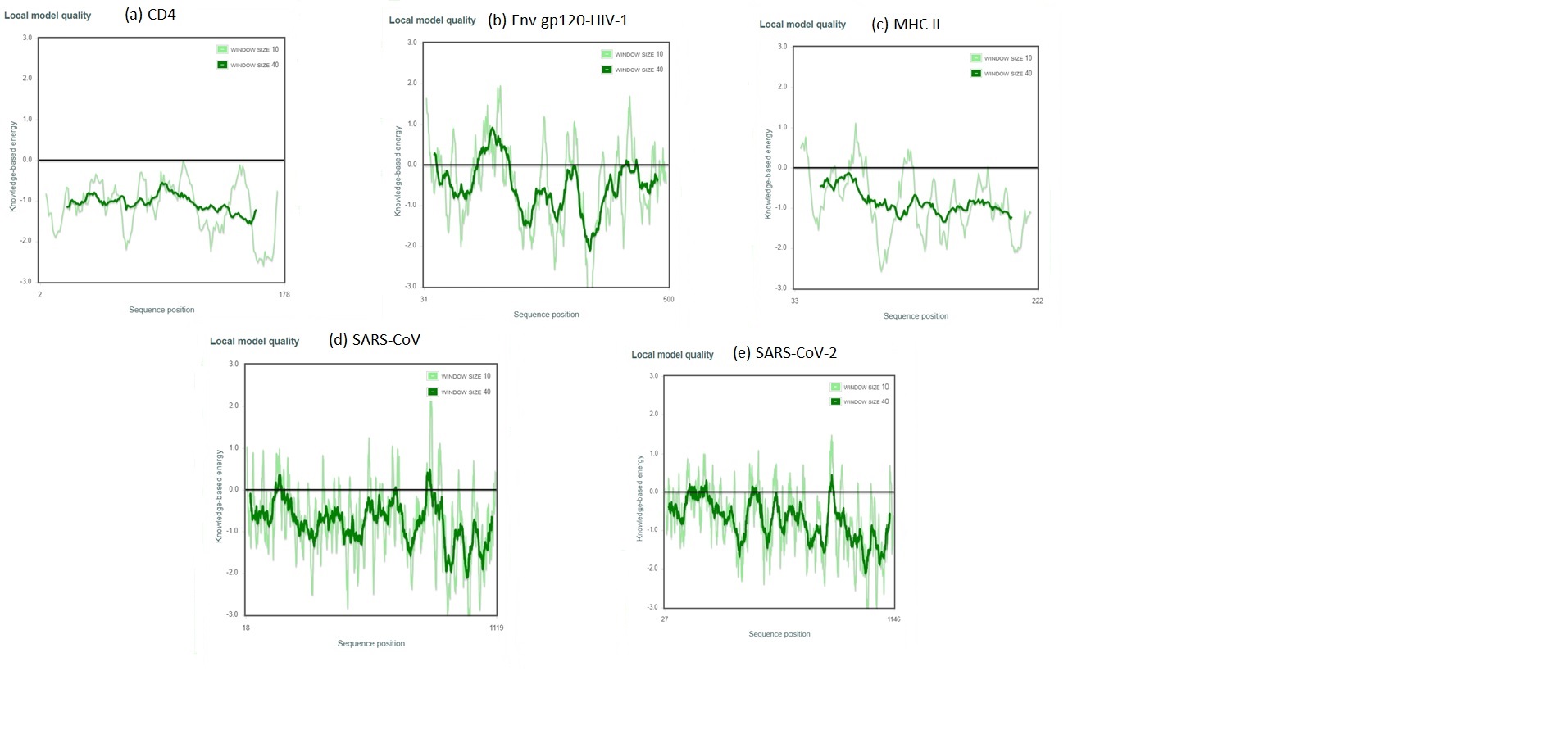
